# Distinct brain network features predict internalizing and externalizing traits in children and adults

**DOI:** 10.1101/2023.05.20.541490

**Authors:** Yueyue Lydia Qu, Jianzhong Chen, Angela Tam, Leon Qi Rong Ooi, Elvisha Dhamala, Carrisa Cocuzza, Connor Lawhead, B. T. Thomas Yeo, Avram J. Holmes

## Abstract

Internalizing and externalizing traits are two distinct classes of behaviors in psychiatry. However, whether shared or unique brain network features predict internalizing and externalizing behaviors in children and adults remain poorly understood. Using a sample of 2262 children from the Adolescent Brain Cognitive Development (ABCD) study and 752 adults from the Human Connectome Project (HCP), we show that network features predicting internalizing and externalizing behavior are, at least in part, dissociable in children, but not in adults. In ABCD children, traits within internalizing and externalizing behavioral categories are predicted by more similar network features concatenated across task and resting states than those between different categories. We did not observe this pattern in HCP adults. Distinct network features predict internalizing and externalizing behaviors in ABCD children and HCP adults. These data reveal shared and unique brain network features accounting for individual variation within broad internalizing and externalizing categories across developmental stages.

## Introduction

A classic distinction in child and adolescent psychiatry has been the study of “internalizing” and “externalizing” behaviors^1^. These two broad classes of psychopathology were first proposed by T.M. Achenbach from a factor analysis of symptoms in children and adolescents with psychiatric illness^2^. Internalizing behaviors are internally directed towards the individual and manifest in their extreme form as sadness, withdrawal, somatic complaints, and anxiety, while externalizing behaviors are directed towards the external environment and involve disruptive, aggressive, impulsive, and defiant behaviors^3^. The expressions of internalizing and externalizing behaviors exhibit cross-generational associations between parents and children^4–6^. These behaviors have also been linked with reduced school engagement and an increased risk for suicide attempts in childhood and adolescence^7–9^, as well as worse work performance and lower cognitive abilities in adulthood^10,11^. However, the neural underpinnings associated with internalizing and externalizing behaviors across distinct developmental stages remain poorly understood.

Throughout development, functional connectivity patterns within and between large-scale brain networks can predict individual differences in cognition^12^, impulsivity^13^ and psychiatric symptoms^14,15^. While individual-level variability in the functioning of large-scale brain networks can predict individual differences within broad categories of cognition, personality and mental health in both children and adults^16,17^, macroscale patterns of brain functioning are dynamic across the lifespan^18–20^. The transition from childhood through adolescence to adulthood reflects critical neurodevelopmental stages characterized by a protracted period of synaptic pruning, intracortical myelination, cortical thinning, and functional network segregation^18,21^. Therefore, it is unclear if the specific brain-behavior relationships observed in childhood mirror those identified in adulthood. Furthermore, although shared network features account for individual variation within broad classes of behavior^16^, individual-specific patterns of functional network connections may predict even finer-grained categories, such as internalizing and externalizing behaviors. Here, we aimed to examine the extent that functional network-based predictors of internalizing versus externalizing behaviors were similar across a large sample of children and their parents. We further tested whether such patterns can be observed in an independent sample of young adults.

In the present study, we predicted internalizing and externalizing measures of psychopathology in a sample of children (and their parents) from the Adolescent Brain Cognitive Development (ABCD) study^22^ using children’s functional connectivity patterns across four brain states: resting-state, monetary incentive delay (MID) task^23^, stop signal task (SST)^24^ and emotional N-Back task^25^. We further explored functional connectivity predictors of internalizing and externalizing behavior in an independent cohort of young adults from the Human Connectome Project (HCP)^26^, using resting-state fMRI connectivity matrices. Multi-kernel ridge regression (multiKRR) models revealed network-based features that were predictive of behaviors within the same category were more correlated with each other than with those across different categories in ABCD children and parents, while KRR models showed a lack of categorical distinction in HCP adults. Moreover, predictive network features were distinct across the two samples. These results support internalizing and externalizing behaviors as distinct factors of psychopathology and suggest that brain-based predictive features may change across the lifespan.

## Results

### ABCD Results

We used fMRI data acquired across three task states, including monetary-incentive delay (MID), stop-signal task (SST) and N-back, as well as resting state fMRI from N=11,875 typically-developing children (ABCD 2.0.1 release^22^). Our analyses considered 33 dimensional measures from the available mental health assessments collected from child participants and their parents^27^, comprised of 15 measures of internalizing problems, 10 measures of externalizing problems, 2 measures of thought problems and 6 measures of attention problems (Supplementary Table 1). The final analytical sample consisted of n=2,262 unrelated children who passed fMRI quality control and had complete data (see Methods).

#### Multi-kernel ridge regression predicts most behavioral measures

We defined 400 cortical and 19 subcortical regions-of-interest (ROIs) based on Schaefer Parcellation^28,29^ and computed a 419 by 419 functional connectivity (FC) matrix for each brain state. Following prior work^16^, we used multi-kernel ridge regression (multiKRR) models to predict each behavioral measure from child-specific FC matrices concatenated across brain states. To evaluate predictive accuracy, we performed nested cross-validation procedures with 120 folds (see Methods). Pearson’s correlation between predicted and actual behavioral scores and coefficient of determination (COD; see Supplementary method S3) were used as accuracy metrics. Statistical significance of prediction accuracy was assessed by permutation testing.

Prediction accuracies -- given by Pearson’s correlation -- of the models trained on children’s functional connectivity data are shown in Fig. 1A (for behavioral predictions in ABCD children) and Fig. 1B (for behavioral predictions in ABCD parents). Most behavioral measures were predicted better than chance after FDR correction (q<0.05), except for child somatic complaints and somatic problems, parent intrusive behavior, parent ADHD problems and parent inattention (see Methods). Parent ADHD problems became significantly predicted after FDR correction when COD was used as the accuracy metric. Prediction accuracies were broadly stable across both metrics (see Supplementary Fig. 1 for COD results). Notably, these findings demonstrate that patterns of FC specific to each child can significantly predict their parent’s self-reported internalizing and externalizing behaviors (Fig. 1B).

**Figure 1.**
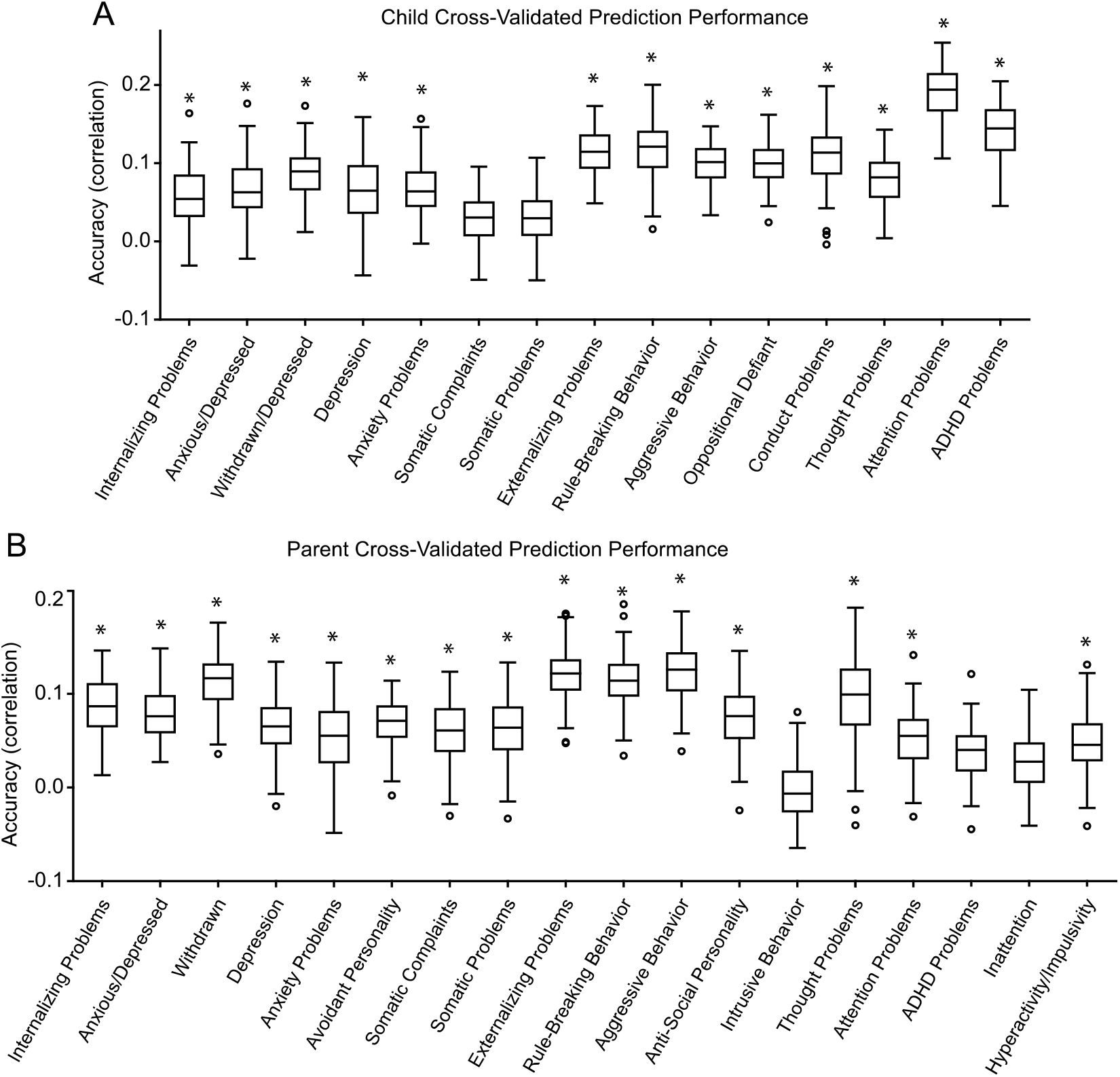
Cross-validated prediction performance using the multi-kernel ridge regression (multiKRR) model, using functional connectivity matrices concatenated across four brain states (resting state, MID, SST and N-back) from children’s neuroimaging data to predict (A) parent-reported child behavior and (B) self-reported parent behavior. Prediction performance was calculated as the mean Pearson’s correlation between observed and predicted values across 120 cross-validation folds for each behavioral measure from the ABCD dataset. For each boxplot, the top and bottom edges represent upper and lower quartiles of correlation coefficient (*r*) distributions, and the horizontal lines mark the corresponding median. Outliers are plotted as circles and were defined as data points outside of the interquartile range. The whiskers extend to the most extreme data points not considered as outliers. Asterisks (*) denote above-chance significance after correcting for multiple comparisons (FDR *q*<0.05).

#### Predictive brain network features are more similar within behavioral categories

There is broad consistency in the brain network features predictive of mental health-relevant traits^16^. Here, we sought to determine if internalizing and externalizing behaviors exhibited unique predictive network markers in childhood. At each cross-validation fold, we quantified “feature importance” (i.e., how important a given network-based predictor was to the model) of each interregional FC edge predicting each behavior using Haufe-transformed (see Methods) predictive feature weights^30^, yielding a 419 by 419 predictive feature matrix for each behavior and for each brain state.

Next, we analyzed whether predictive feature weights computed from multiKRR model outputs were more similar among behaviors within than between categories (Fig. 2). The predictive feature weight vector for each behavioral measure was averaged across all four brain states and correlated with all other measures. Focusing on each of the four internalizing and externalizing categories (Child Internalizing, Child Externalizing, Parent Internalizing and Parent Externalizing), the difference between mean correlation within each category (“within-category mean correlation”) and mean correlation with all other three categories (“between-category mean correlation”) was computed 10000 times and used to generate a null distribution of mean differences (Fig. 3; see Methods). Mean within-category correlations of predictive feature weights were significantly higher than mean between-category correlations (FDR *q*s≤0.002; Fig. 3). The above analyses were rerun using KRR models using only resting-state fMRI and yielded similar results (Supplementary Fig. 2-3). Notably, the similarity pattern of predictive feature weights across behavioral measures was highly correlated with the similarity pattern of these behavioral measures on the behavioral level (Supplementary Fig. 4; *r*=0.97).

**Figure 2.**
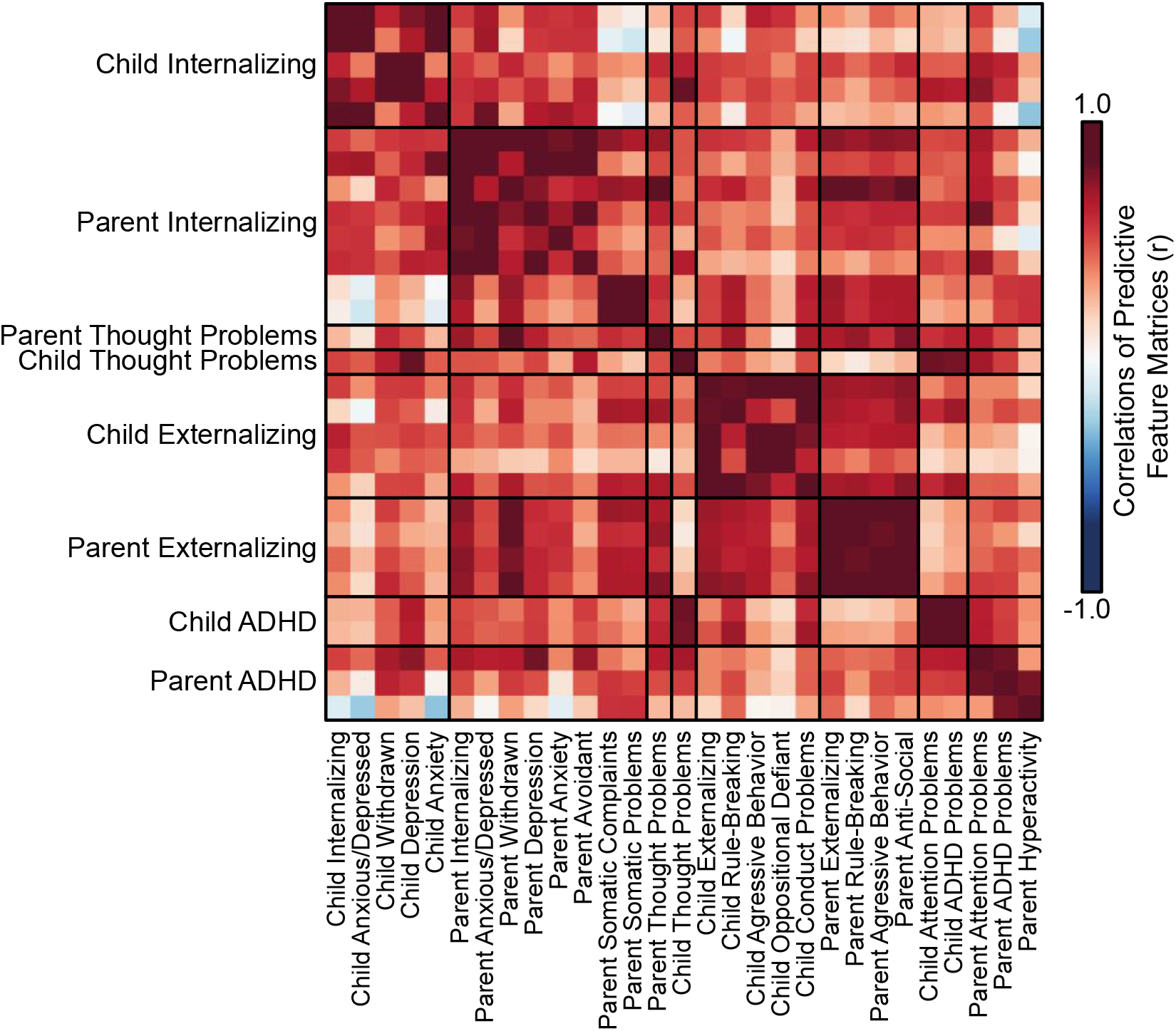
Predictive-network features are similar within behavioral categories. Pearson’s correlation (*r*) of predictive feature weights between all pairs of behavioral measures significantly predicted by multiKRR models in the ABCD study. Behavioral measures from the same behavioral categories are grouped together. Warmed colors indicate stronger positive correlations of the mean predictive feature weights between a pair of behavioral measures, indicating that these behavioral measures were predicted by similar functional connectivity patterns.

**Figure 3.**
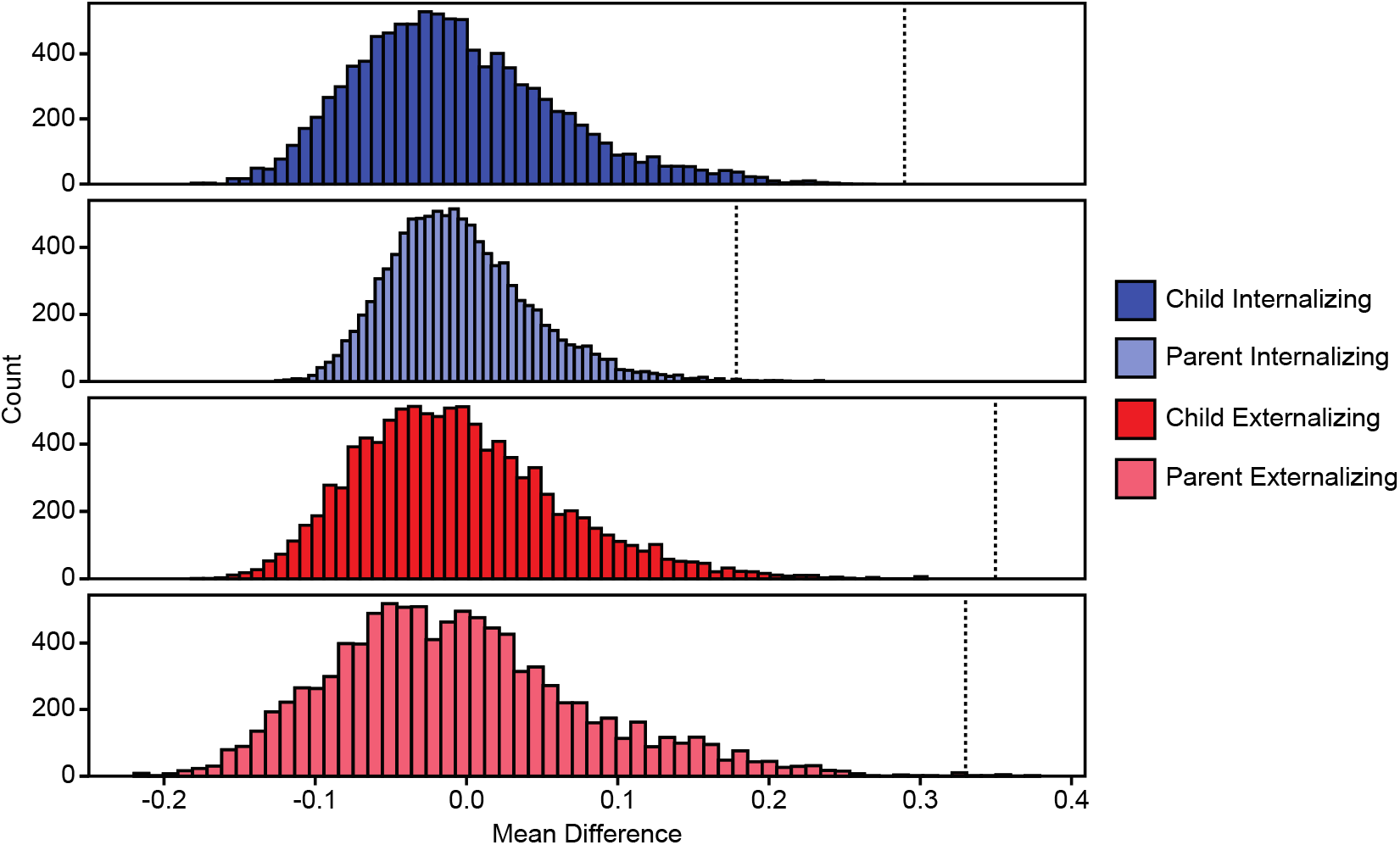
Correlations of predictive feature weights computed from multiKRR outputs were significantly stronger across behavioral measures within the same category than between different categories. Differences between within- and between-category mean correlations for child and parent internalizing and externalizing categories were significantly greater than the null distributions (FDR *qs* ≤0.002). Correlation values were converted to z-scores using Fisher’s r-to-z transformation prior to averaging. Histograms display null distributions of mean differences generated through 10000 permutations with shuffled behavioral labels. Dashed lines represent observed mean differences for each of the four categories.

#### Distinct brain network features in children predict internalizing and externalizing behaviors in both children and their parents

Prior work suggests the presence of shared network features across broad categories of mental health^16^. Our analyses revealed unique parcel-level FC profiles predicting distinct aspects of psychopathology. Next, we examined the extent that some predictive-network features may be shared across behavioral categories. Predictive feature matrices were averaged across all behavioral measures within each category, resulting in 32 predictive feature matrices (one for each behavioral category and each brain state). To limit the number of multiple comparisons, predictive feature weights were averaged within and between 18 networks (following the 17-network partition in Yeo et al., 2011^31^ plus one subcortical network^28^) at each permutation. Permutation testing was performed on mean predictive feature weights from each of the resulting 171 unique network blocks. We also conducted a conjunction analysis to extract the predictive feature weights that were not only statistically significant but also exhibited consistent directionality (positive or negative) across all brain states, and then averaged these predictive feature weights across all brain states (Fig. 4A). These analyses yielded predictive feature weights that are both shared across behavioral measures within a category and across brain states (Fig. 4B). Predictive feature weights were summed across each row in Fig. 4A and plotted on brain surface in Fig. 4C for the positive weights and Fig. 4D for the negative weights. These figures reveal that both shared and unique FC patterns predict distinct behavioral categories in both children and their parents.

**Figure 4.**
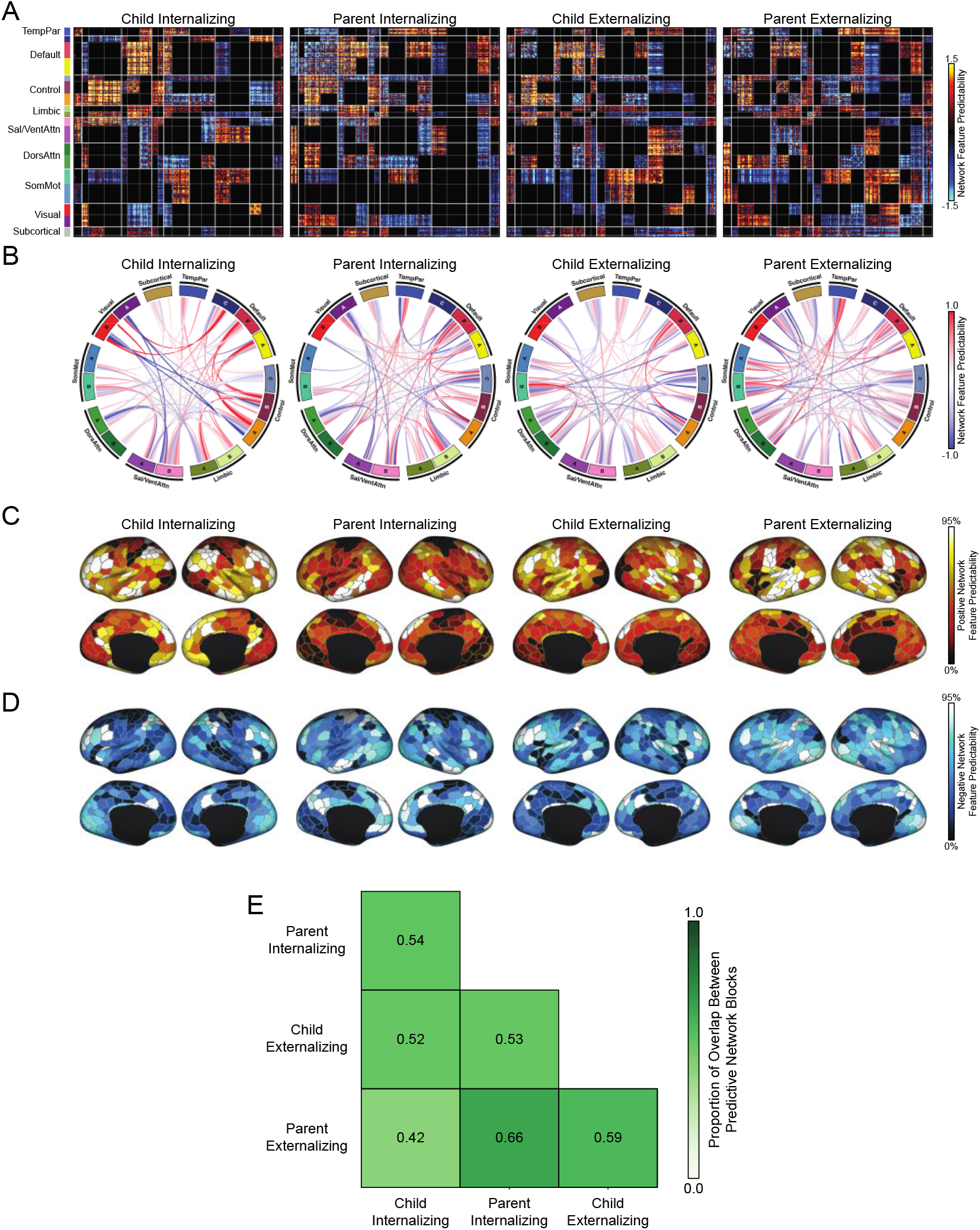
Shared and unique functional network features predict internalizing and externalizing behaviors in children and their parents. (A) Matrices of predictive feature weights, averaged across all behavioral measures within each child and parental internalizing and externalizing categories, and averaged across all brain states. Only weights that were statistically significant and that exhibited the same directionality across all brain states are shown. Rows and columns: predictive weights based on FC estimates of all pairwise cortical regions. For visualization purposes, all predictive feature weights were divided by their standard deviation. (B) Predictive feature weights averaged based on network assignment in panel (A). (C) Positive predictive feature weights summed across rows of panel (A) for each cortical region. A more positive value indicates that stronger functional connectivity associated with a given cortical parcel predicts higher behavioral scores in a behavioral category. (D) Negative predictive feature weights summed across rows of panel (A) for each cortical region. A more negative value indicates that weaker functional connectivity associated with a given cortical region predicts higher behavioral scores in a behavioral category. In both panels (C) and (D), the color of each cortical region indicates the percentile of predictive feature weights among 400 regions. (E) The 2D grid displays the proportion of network blocks that exhibit the same directionality across each pair of child behavioral categories relative to the behavioral category represented by each column. Here, each within- and between-network block was coded as 1, 0 or -1 depending on whether sum of predictive feature weights within that block is greater than, equal to or lesser than 0, resulting in an 18 by 18 matrix for each behavioral category. The number of network blocks having the same non-zero entries across both matrices associated with each pair of behavioral categories was counted and divided by the total number of non-zero significantly predictive network blocks.

To examine the extent to which these predictive features are similar across behavioral categories in children and their parents, we next calculated the proportion of overlapping network blocks which significantly predicted each pair of behaviors (Fig. 4E). Two network blocks were counted as overlapping if sums of predictive feature weights within these network blocks exhibited consistent directionality. Of note, the observed predictive features were not fully distinct across children and parents. The largest proportion of overlap was 0.66 between parent internalizing and externalizing categories, while the lowest proportion of overlap was 0.42 between child internalizing and parent externalizing categories. Proportions of overlap between other four category pairs ranged from 0.50 to 0.60, demonstrating the presence of both common and distinct patterns of predictive-network features across categories. As one example, the proportion of network blocks that exhibit the same directionality across child and parent internalizing categories was 0.54.

#### Single-kernel ridge regression predicted most behavioral measures in adults

To examine brain-based predictive network features in adults, we used resting-state fMRI data from the HCP WU-Minn S1200 sample (n=752) and analyzed 18 dimensional measures from the Achenbach Self-Report^32^. There were 8 measures of internalizing problems, 5 measures of externalizing problems, 1 measure of thought problems and 4 measures of attention problems (Supplementary Table 2; see Methods). All analysis steps were performed as above, except that single-kernel ridge regression (KRR) models were used to predict each behavioral measure from subject-specific resting-state FC due to the lack of task fMRI data in the HCP. Given that the HCP was not collected across different sites, we implemented 60 random initiations of 10-fold nested cross-validation.

Prediction accuracies – given by Pearson’s correlation – of the KRR models across all behaviors are shown in Supplementary Fig. 6. Although most behavioral measures were predicted better than chance after FDR correction (q<0.05), only two out of eight behavioral measures under the internalizing category survived FDR correction (Supplementary Fig. 6). When COD was used as the accuracy measure, only withdrawn, aggressive behavior, and attention problems reached better-than-chance accuracy after FDR correction (Supplementary Fig. 7).

#### Predictive brain network features are similar across behavioral categories in adulthood

As above, we examined similarity patterns of predictive feature weights calculated from KRR model outputs across behaviors within and between different categories (Fig. 5). In contrast to the ABCD analyses, predictive feature weights were highly correlated across categories (Fig. 5; Supplementary Fig. 9). We then conducted a permutation test similar to the ABCD analyses, focusing on adult internalizing and externalizing categories. Mean within-category correlations of predictive feature weights were not significantly different from mean between-category correlations (FDR *qs*>0.12; Fig. 5B). These results suggest that predictive network features associated with internalizing and externalizing behavior in adults are broadly consistent between behavioral categories. Although network features predicting intrusive behavior were weakly correlated with those predicting other measures, it is not surprising given the weak correlations between intrusive behavior and other measures on the behavioral level. The observed similarity pattern of predictive feature weights across behaviors was moderately correlated with the similarity pattern of these measures on the behavioral level (Supplementary Fig. 8; r=0.59).

**Figure 5.**
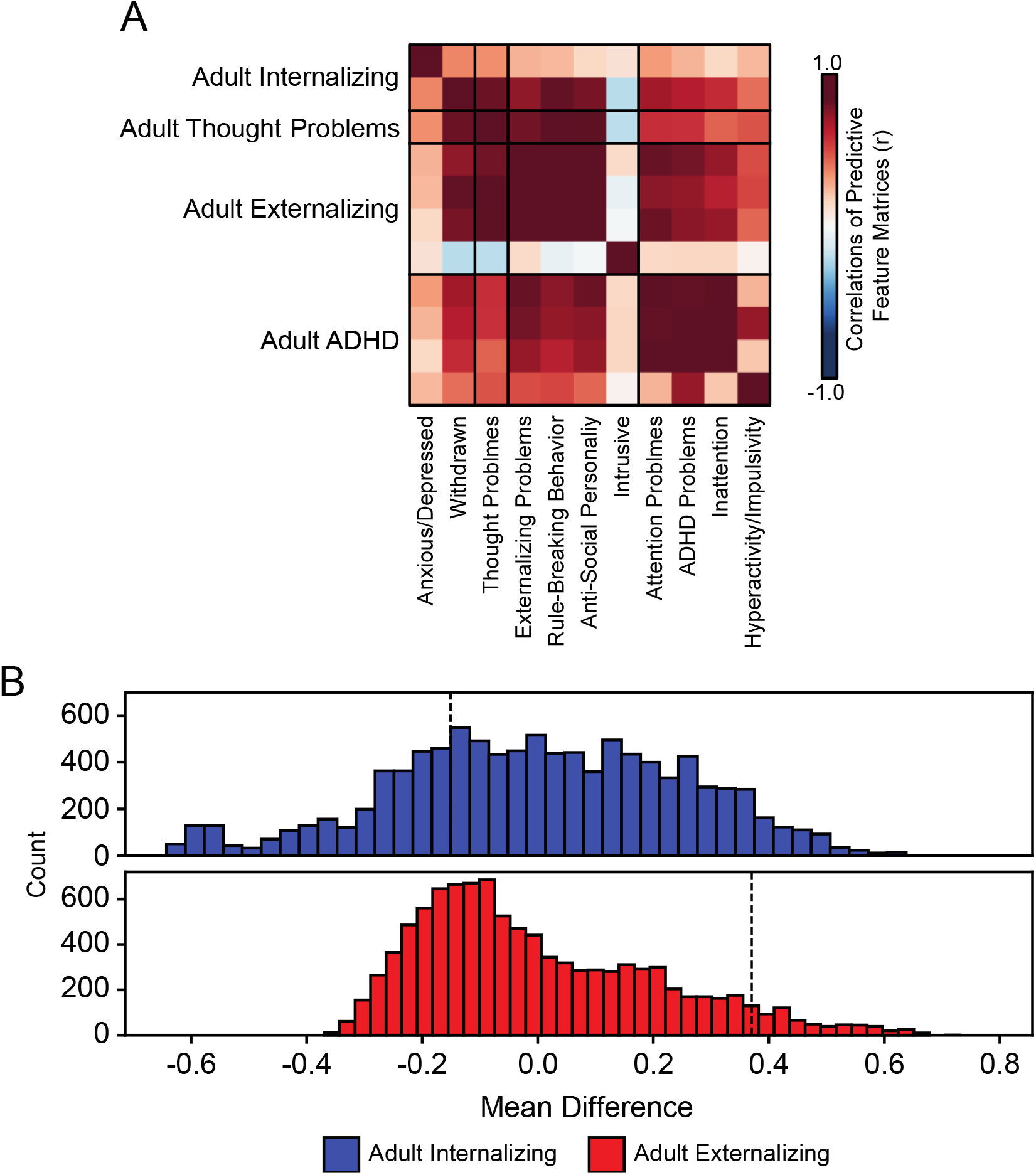
Predictive brain network features are similar across conceptually-linked behavioral categories in adulthood. (A) The correlation matrix displays Pearson’s correlation *r* of predictive feature weights between all pairs of behavioral measures significantly predicted in the HCP study. Behavioral measures are grouped within associated behavioral categories. Higher intensity colors indicate higher positive (red) and negative (blue) correlations of the mean predictive feature weights between a pair of behavioral measures. (B) Differences between mean correlations of predictive feature weights across behavioral measures within the same category and between different categories were not significantly greater than the null distributions in the adult internalizing and externalizing categories (FDR *qs*>0.12). Correlation values were converted to z-scores using Fisher’s r-to-z transformation prior to averaging. Graphing conventions are similar to that of Figure 3.

#### Predictive feature weights are largely distinct across ABCD and HCP datasets

To investigate if functional network features predicting internalizing and externalizing behaviors are distinct across development, we assessed the similarity of predictive network features computed from KRR model outputs between ABCD and HCP datasets. We found that FC features predicting internalizing and externalizing problems in ABCD children and HCP adults were only weakly correlated (Fig. 6). We then ran permutation tests to compare the difference in the mean correlation within each category and the mean correlation between each category and the corresponding category in the other age group. As the adult internalizing category contained only two significantly predicted measures, we only focused on child internalizing and externalizing and adult externalizing behaviors. The difference was significantly greater than its null distribution (FDR *qs*≤0.0146) for the two child categories but failed to reach statistical significance the adult externalizing category (FDR *q*=0.0601; Fig. 6B). From the predictive feature matrices associated with child and adult internalizing and externalizing categories, we computed the proportion of overlapping network blocks which significantly predicted each pair of categories (Fig. 6C). Proportions of overlap were distinctly higher for pairs of behavioral categories within the same dataset than between the two datasets. These results suggest that although shared brain network features account for individual variation within broad categories of internalizing and externalizing problems in childhood, functional network predictors may change throughout the lifespan, exhibiting distinct fingerprints across developmental stages.

**Figure 6.**
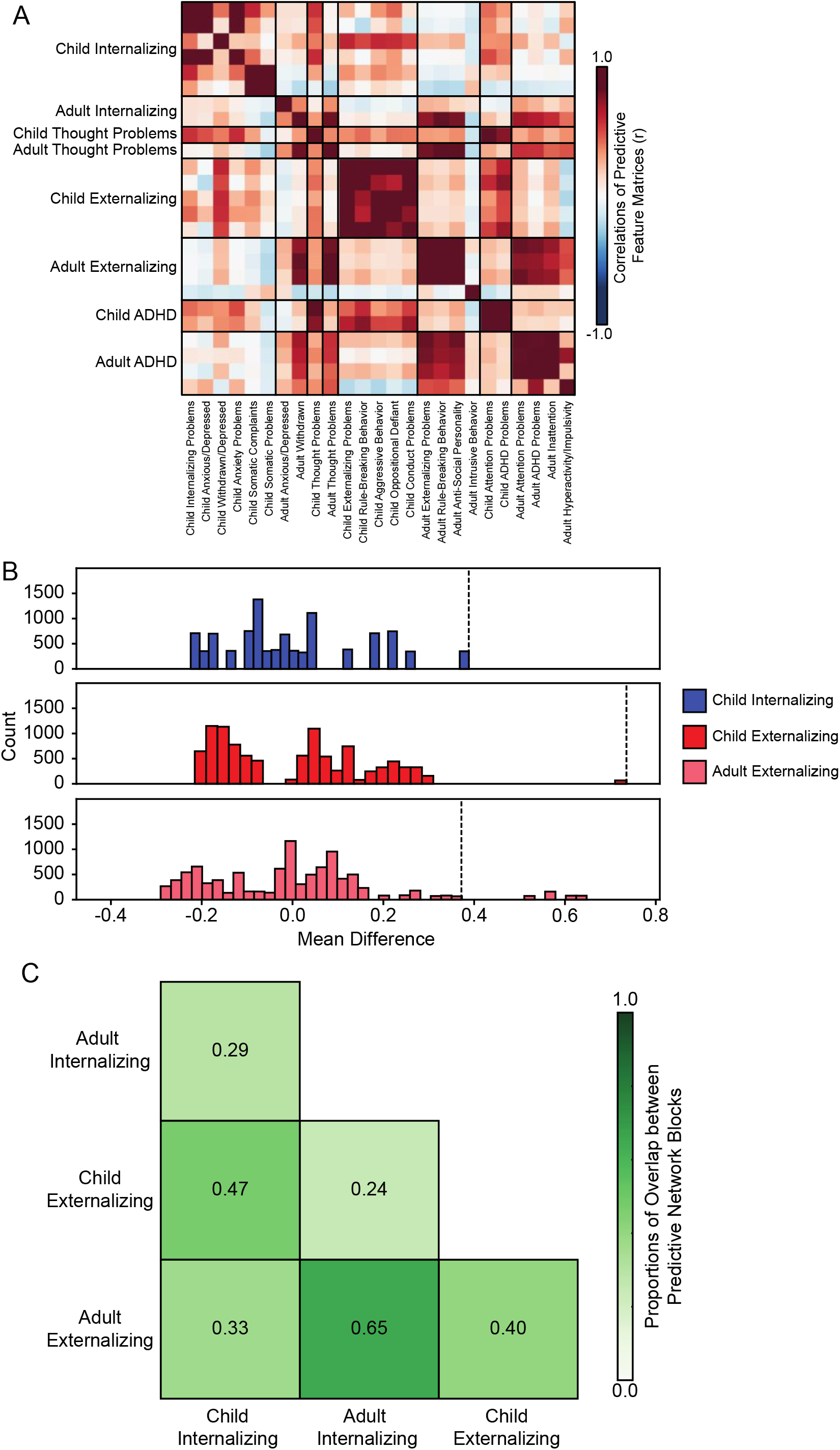
Distinct functional network features predict internalizing and externalizing behaviors in children from the ABCD study and adults from the HCP study. (A) The correlation matrix displays the Pearson’s correlation *r* of predictive feature weights between all pairs of behavioral measures significantly predicted by KRR models in the ABCD and HCP studies. Measures from the same behavioral category are grouped together. Colors indicate positive (red) and negative (blue) correlations of the mean predictive feature weights between behavioral measures and populations. (B) Predictive feature weights associated with child internalizing and externalizing categories were significantly more correlated within categories than with the corresponding adult categories (FDR *qs*≤0.0186). The permutation test was not applicable for the adult internalizing category because only one correlation can be computed between two behavioral measures in the category. Correlation values were converted to z-scores using Fisher’s r-to-z transformation prior to averaging. Graphing conventions are similar to that of Figure 3. (C) The 2D grid displays the proportion of network blocks that exhibit the same directionality across each pair of child and adult behavioral categories relative to the behavioral category represented by each column.

Despite the broad distinction between FC patterns predicting internalizing and externalizing behaviors across these two datasets, shared predictive patterns may still be present within select edges. Here, the shared predictive network features associated with both child and adult internalizing categories primarily involved the default, control and visual networks, while the shared predictive network features associated with child and adult externalizing primarily involved somato/motor, ventral and dorsal attention networks (Supplementary Fig. 10).

## Discussion

In this study, we first used estimates of individual-specific, functional connectivity from a large, diverse sample of healthy children to predict internalizing- and externalizing-related behavioral measures in children and their parents. Predictive feature weights were more correlated across behaviors within the same categories than with those from different categories. Of note, our analyses revealed that brain data specific to each child can be used to predict self-reported internalizing and externalizing behaviors in their parents. We repeated these analyses in an independent sample of healthy young adults and the opposite pattern was observed, where predictive feature weights were similarly correlated across distinct mental health-linked behavioral categories. Moreover, predictive feature weights associated with internalizing- and externalizing-related behavioral measures were distinct across children and adults, suggesting that brain-based predictors of internalizing and externalizing behaviors may change across the lifespan.

Internalizing and externalizing symptoms reflect distinct factors across various mental disorders, irrespective of demographic and collection method^33–37^. Predictive network features are similar across behaviors within the broad categories of mental health^16^. Although large-scale networks can be mechanistically informative for studying neurocognitive processes^38,39^ and psychiatric phenotypes^15,40–42^, the similarity of whole-brain FC patterns predicting measures of internalizing and externalizing behavior has not been directly assessed. Through the use of both KRR and multiKRR models^16,43^, we were able to predict most mental health measures in children and their parents from children’s resting-state functional connectome. Here, we demonstrated that the whole-brain patterns of functional connectivity in children can be used to predict internalizing and externalizing measures in their parents. Our results highlight that the predictive utility of functional connectomes may extend beyond the individual, and provide a robust entry point for future work on shared environmental and contextual factors, broader behavioral patterns within family systems, and/or the heritability of internalizing and externalizing traits.

Consistent with prior work by Chen et al. 2022 (which used the same dataset), we observed that predictive features are generally similar across measures of internalizing and externalizing behaviors. However, above and beyond this broad pattern of similarity, predictive feature weights were more correlated within than between behavioral behavioral categories in ABCD children. These findings are consistent with theoretical models that consider internalizing and externalizing behaviors as distinct constructs of psychopathology under a general psychopathology *p* factor^44,45^. Behavioral measures associated with different categories are characterized by both common and distinct network predictors in children. On average, higher behavioral scores in both child internalizing and externalizing categories were predicted by more positive FC between default, control and limbic networks, between somato/motor and salience networks and more negative FC between default and somato/motor networks. Beyond these shared features, there was substantial heterogeneity in the FC patterns predicting internalizing- and externalizing-related behaviors. These results align with previous neuroimaging studies implicating frontoparietal^46,47^, default^47–49^, salience^49,50^, limbic^49^ and somato/motor^49,51^ network disruptions across psychiatric disorders.

Contrary to the similarity pattern observed in ABCD children, mean correlations of predictive feature weights across all pairs of behavioral measures within internalizing and externalizing categories were not significantly different from mean correlations between different categories in HCP adults. Our findings suggest that diffuse functional network patterns may predict a more general psychopathology factor in adults, while more specific FC patterns may differentially predict behaviors associated with specific categories of psychopathology in children. One consideration is that KRR models used in HCP analyses reached better-than-chance predictive accuracy for only two out of eight measures assigned to adult internalizing category. This may have biased the results for the permutation test assessing statistical significance of mean correlation differences within and between adult internalizing and externalizing categories. In addition to different correlation patterns across behavioral categories between the two datasets, we also observed distinct FC features predicting same categories of internalizing and externalizing behaviors in children compared to adults. Weak correlations of predictive feature weights associated with internalizing and externalizing behavior across the two samples may be attributable to development of functional network organization from childhood through adolescence and then adulthood^19,20,52–54^. However, such differences may also be attributable to site differences between the two collection efforts. Of note, our interpretations are limited by the cross-sectional nature of the available data. Future work should further characterize the longitudinal trajectories of brain development and associated brain-based predictions across the lifespan. Another limitation of our study is that we did not test our models separately in each sex. Previous studies have suggested brain-based predictive models often fail to generalize across sexes^55^, and future work should test sex-/gender-specific models of behavior^56^.

Taken together, our study found that predictive network features cluster within the same categories of internalizing and externalizing behavior in ABCD children. Intriguingly, the utility of brain-based predictive models in children extended to capture behaviorally relevant signals in their parents. Finally, although most behaviors were predicted better than chance in children and adults, analyses revealed distinct network predictors across datasets. Future work will benefit from the longitudinal study of common and distinct brain-based predictive features across childhood, adolescence, and adulthood.

## Methods

### Participants

11,875 typically-developing children and their parents across 21 sites in the United States participated in the ABCD study at baseline (ABCD release 2.0.1). The final analytical sample consisted of 2,262 unrelated children who passed strict preprocessing quality control, had complete fMRI data across all brain states and complete scores across all behavioral measures. Similar to Chen et al., 2022, we combined the 22 ABCD sites into 10 “site-categories” to reduce sample size variability across sites (Supplementary Table 5). Subjects within the same site were also in the same site-category. Detailed demographic information can be found in Supplementary Table 6.

1,206 healthy adults participated in the HCP study (HCP S1200 Data Release). After pre-processing quality control of imaging data, participants were filtered from Li’s set of 953 participants^57^ based on the availability of a complete set of structural and resting-state fMRI scans, as well as all behavioral scores of interest. Our main analysis comprised 752 adult participants, who fulfilled all selection criteria^17^. Detailed demographic information can be found in Supplementary Table 7.

### Neuroimaging

#### Data acquisition

For the ABCD study, all T1w images and fMRI data was acquired using protocols harmonized across three 3 tesla(T) scanner platforms (i.e., Phillips, Siemens Prisma and General Electric 750) at 21 sites. Twenty minutes of resting-state fMRI data, consisting of four 5-minute runs, was collected from each ABCD child participant. For each of the three tasks (MID, SST and N-Back)^23–25^, fMRI data was acquired over two runs with 2.4mm isotropic resolution with a TR of 800ms. The structural T1 scans were acquired with 1mm isotropic resolution with a TR of 2500ms. For full details of imaging acquisition can be found elsewhere^58^.

The fMRI data in the HCP data was acquired using an optimized protocol with 2mm isotropic resolution and a TR of 700ms. Each HCP subject goes through one structural MRI session and two fMRI sessions. Each fMRI session consists of two 15-minute resting-state scans with opposite phase encoding directions (L/R and R/L). The structural T1 scans were acquired using 0.7mm isotropic resolution and a TR of 2400ms. Full details of the acquisition protocol can be found elsewhere^26^.

#### Data processing

Minimally preprocessed T1w images^59^ in the ABCD study were further processed using FreeSurfer v5.3.0^60–65^. The cortical surface meshes were then registered a common spherical coordinate system^62,63^. Subjects who failed recon-all QC were subsequently excluded^59^. The minimally preprocessed fMRI data^59^ were subsequently processed in the following manner. The first four frames were removed^59^. Slice time correction was performed with the FSL library^66,67^. Motion correction was performed using rigid body translation and rotation with the FSL package. The resulting fMRI images were then aligned with the processed T1w images^68^ with FsFast (https://surfer.nmr.mgh.harvard.edu/fswiki/FsFast), and only runs with registration costs less than 0.6 were retained. Framewise displacement (FD)^67^ and voxel-wise differentiated signal variance (DVARS)^69^ were computed by fsl_motion_outliers. Volumes with FD > 0.3 mm or DVARS > 50, along with one volume before and two volumes after, were flagged as outliers. A bandstop filter was applied to remove respiratory pseudomotion^70^. Uncensored segments of data having fewer than 5 contiguous volumes were also flagged as outliers and censored^71^. Runs with more than half of the volumes flagged as outliers were discarded. Participants with less than 4 minutes of data for each fMRI state (rest, MID, N-Back, SST) were excluded from further analysis. Nuisance regressors, including global signal, six motion correction parameters, averaged ventricular signal, averaged white matter signal, and their temporal derivatives (18 regressors in total), were regressed out of the fMRI time series from the unflagged volumes. Data were interpolated across censored frames^72^, band-pass filtered at 0.009 Hz≤f≤0.08 Hz, projected onto FreeSurfer fsaverage6 surface space, and smoothed using a 6mm full-width half maximum kernel.

For the HCP study, minimally preprocessed T1w images^73^ went through bias- and distortion-correction using the *PreFreeSurfer* pipeline and registered to MNI space. Cortical surface reconstruction was conducted using FreeSurfer v5.2 using recon-all adapted for high-resolution images. The reconstructed surface meshes were then registered to the Conte69 surface template^74^. After preprocessing, the fMRI data were corrected for gradient-nonlinearity-induced distortions. The fMRI time series in each frame were then realigned to the single-band reference image to correct for subject motion using rigid body transformation^67,75^ with FSL. The resulting single-band image underwent spline interpolation to correct for distortions and was then registered to the T1w image^68^. Native fMRI volumes went through nonlinear registration to the MNI space and mapped to the standard CIFTI grayordinate coordinate space. Further details about the preprocessing and processing pipelines of structural and functional images can be found elsewhere^73^.

#### Functional connectivity

We used 400 cortical regions of interest^29^ (ROIs) and 19 subcortical ROIs^28^. Functional connectivity (FC) was measured by Pearson’s *r* correlations between the mean time series of each pair of ROIs. Censored frames were ignored when computing functional connectivity. In the ABCD study, the average FC matrix across all runs in each subject from each state (rest, MID, N-back, SST) was used for subsequent analyses. To match processing across resting and task states, task activations were not regressed from the task-state data. For the HCP study, the average FC matrix across all runs in each subject was only computed from the resting state and used for subsequent analyses.

### Measures of internalizing and externalizing behaviors

We included 25 dimensional measures of internalizing and externalizing in our analyses, selected from all available mental health relevant assessments taken from child participants and their parents^27^ in the ABCD study. This consisted of 15 internalizing measures and 10 externalizing measures. Thought and attention problems are related to both internalizing and externalizing psychopathology^76,77^. Accordingly, we included 2 measures of thought problems and 6 measures of attention. Participants without available data across all behavioral measures were excluded from analysis. The complete list of the included variables can be found in Tables S1 and S2. Behavioral measures were grouped into four categories: Internalizing, Externalizing, Thought Problems and ADHD Problems for both children and their parents, resulting in eight behavioral categories in total (Supplementary Table 1).

In data from the HCP, we analyzed 18 dimensional measures of internalizing, externalizing, thought and attention problems from the Achenbach Self-Report (ASR) questionnaire, resulting in four behavioral categories (Supplementary Table 2).

### Statistical analysis

Consistent with prior work^16^, we used multi-kernel ridge regression (multiKRR) with l_2_ regularization to predict each behavioral measure from participant-specific FC matrices across all brain states (rest, MID, N-back, SST) jointly in the ABCD study. Behavioral measures in the HCP study were predicted from resting-state FC using kernel ridge regression (KRR) with l_2_ regularization. Details about KRR and multiKRR models can be found in the Supplement (see Supplementary methods S1 and S2). Age and mean FD were entered as covariates. Both models assume that participants with more similar FC patterns have more similar behavioral measures. Models were implemented with nested cross-validation procedures similar to Ooi et al., 2022. Head motion (mean FD and DVARS) were regressed from each behavioral measure prior to cross-validation.

In the ABCD analyses, we performed leave-3-site-clusters-out nested cross-validation for each behavioral measure. At each fold, a different set of 3 site-categories served as the test set, and the remaining 7 site-categories were used as the training set, resulting in 120 folds in total. In the HCP analyses, we implemented 60 random initiations of 10-fold nested cross-validation. Participants from the same family were assigned to either training or testing sets and were never split across training and test sets in any cross-validation fold.

Across both datasets, model and regularization parameters were estimated from the training set at each fold. The estimated parameters were then applied to the unseen participants from the test set and evaluated for accuracy by both correlating predicted and actual measures^78^, and by coefficient of determination (COD). To assess whether model prediction performed better than chance, statistical significance of prediction accuracy was assessed by a permutation test whereby the entire cross-validation procedure was rerun on behavior measures randomly reshuffled across participants in each dataset. Care was taken to avoid shuffling between families or sites.

### Model Interpretation

To interpret the predictive importance of each FC feature, we used an approach from Haufe and colleagues (2014) to transform predictive feature weights associating each FC edge to the behavioral measure. Predictive feature weight was computed by the covariance between the predicted behavioral measure and the FC edge. This resulted in a 419 x 419 predictive feature matrix for each brain state and each behavioral measure. A positive (or negative) predictive feature weight indicates that higher FC predicts greater (or lower) behavioral values. Statistical significance of these predictive feature weights was tested with permutation tests and corrected for multiple comparison using FDR (q<0.05). To reduce the number of multiple comparisons, predictive feature weights were averaged within and between 18 large-scale functional networks^28,29^ before conducting the permutation test.

## Acknowledgements

Data used in the preparation of this article were obtained, in part, from the Adolescent Brain Cognitive Development ^SM^ (ABCD) Study (https://abcdstudy.org), held in the NIMH Data Archive (NDA). This is a multisite, longitudinal study designed to recruit more than 10,000 children aged 9-10 and follow them over 10 years into early adulthood. The ABCD Study® is supported by the National Institutes of Health and additional federal partners under award numbers U01DA041048, U01DA050989, U01DA051016, U01DA041022, U01DA051018, U01DA051037, U01DA050987, U01DA041174, U01DA041106, U01DA041117, U01DA041028, U01DA041134, U01DA050988, U01DA051039, U01DA041156, U01DA041025, U01DA041120, U01DA051038, U01DA041148, U01DA041093, U01DA041089, U24DA041123, U24DA041147. A full list of supporters is available at abcdstudy.org/federal-partners.html. A list of participating sites and a complete list of the study investigators can be found at abcdstudy.org/consortium_members/. ABCD consortium investigators designed and implemented the study and/or provided data but did not necessarily participate in the analysis or writing of this report. Additional data were provided, in part, by the Human Connectome Project, WU-Minn Consortium (Principal Investigators: David Van Essen and Kamil Ugurbil; 1U54MH091657) funded by the 16 NIH Institutes and Centers that support the NIH Blueprint for Neuroscience Research; and by the McDonnell Center for Systems Neuroscience at Washington University. This manuscript reflects the views of the authors and may not reflect the opinions or views of the NIH or the ABCD and HCP consortia investigators.

