## Supplementary Information for "Distinct brain network features predict internalizing and externalizing traits in children and adults"

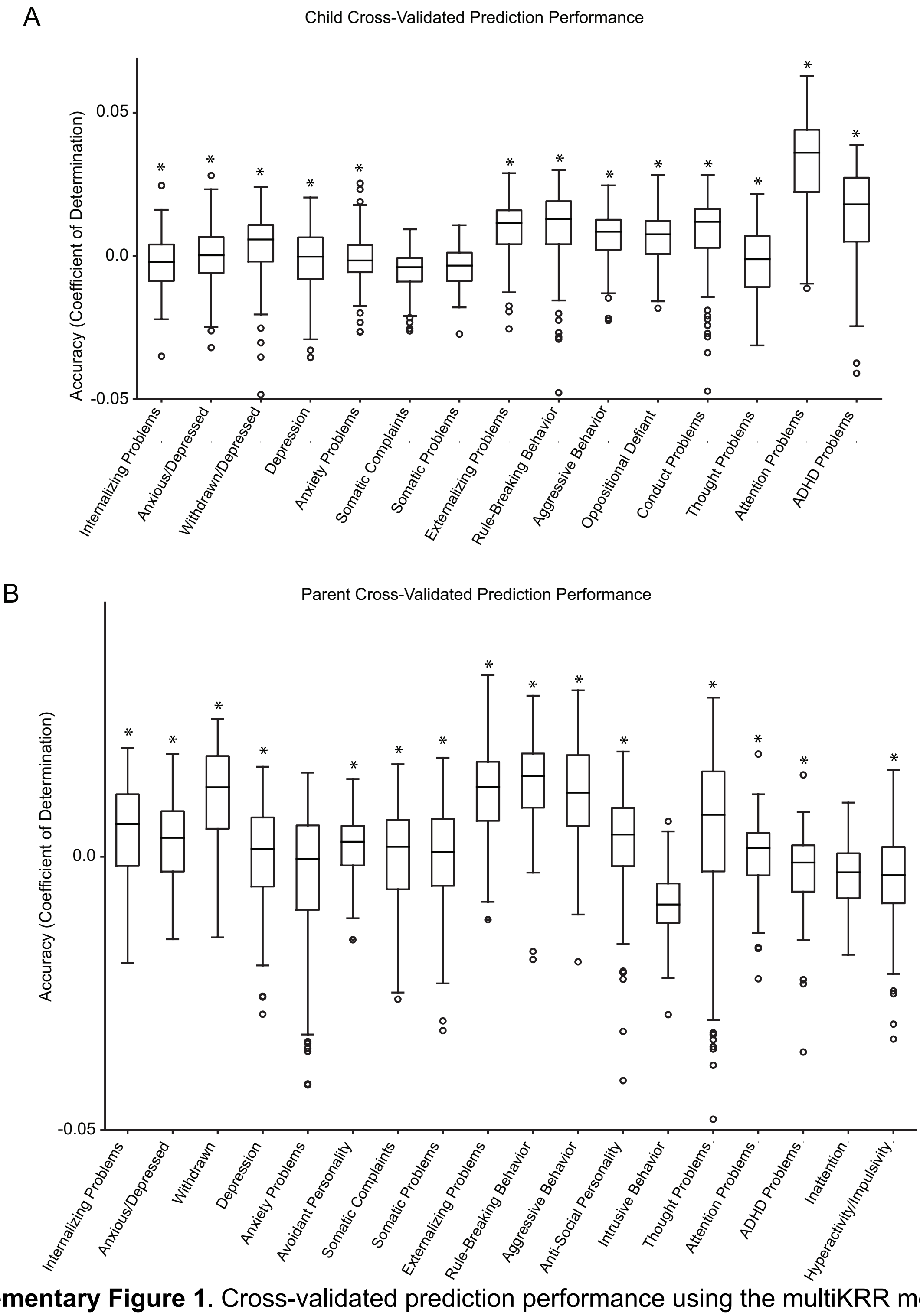


**Supplementary Figure 1**. Cross-validated prediction performance using the multiKRR model applied to functional connectivity matrices concatenated across four brain states (resting state, MID, SST and N-back), in children applied to self-reported (A) child and (B) parent behavior. Prediction performance is calculated as mean COD across 120 cross-validation folds for each behavioral measure from ABCD. For each boxplot, the top and bottom edges represent upper and lower quartiles of COD distribution, and the horizontal line marks the median. Outliers are plotted as circles and are defined as data points outside of the interquartile range. The whiskers extend to the most extreme data points not considered outliers. *s denote above-chance significance after correction for multiple comparisons (FDR $q<0.05$).

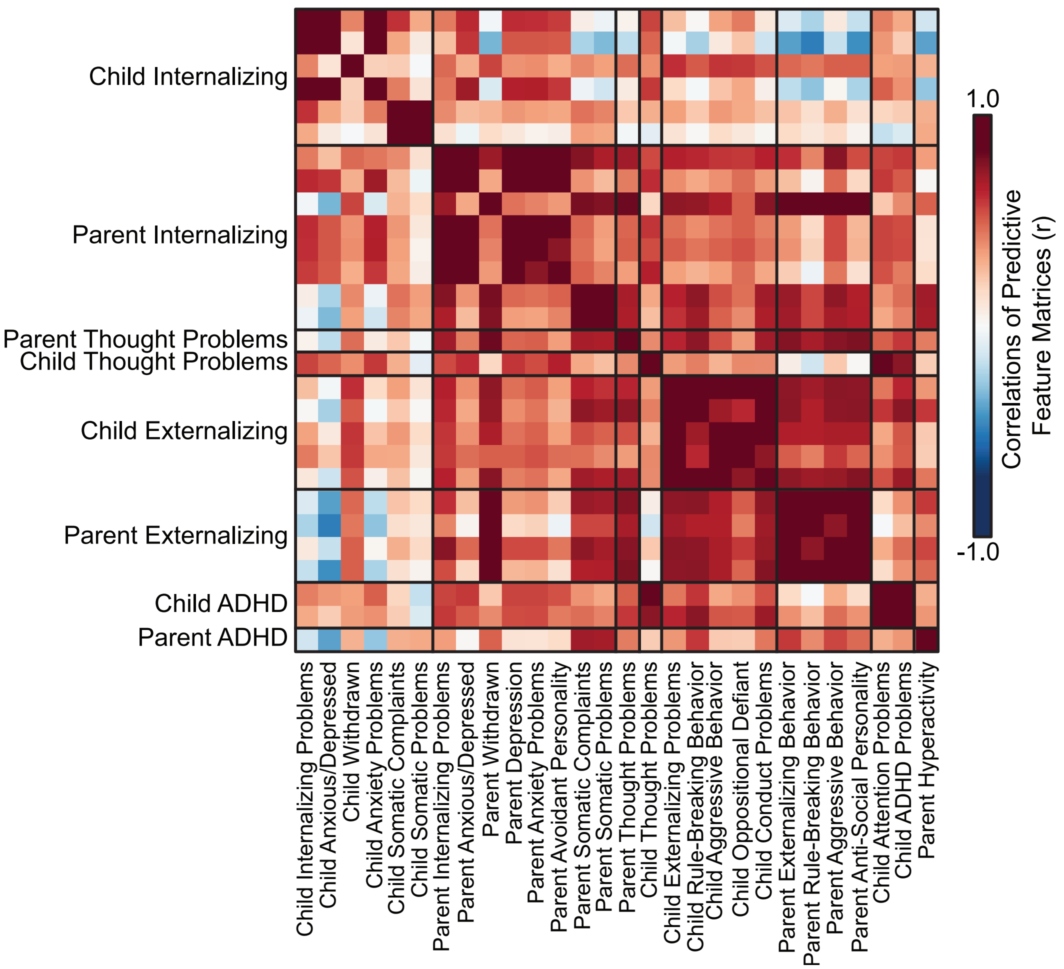


**Supplementary Figure 2.** Predictive-network features are similar within behavioral categories. Pearson's correlation (*r*) of predictive feature weights between all pairs of behavioral measures significantly predicted by KRR models in the ABCD study. Behavioral measures from the same behavioral categories are grouped together. Warmed colors indicate stronger positive correlations of the mean predictive feature weights between a pair of behavioral measures, indicating that these behavioral measures were predicted by similar functional connectivity patterns.

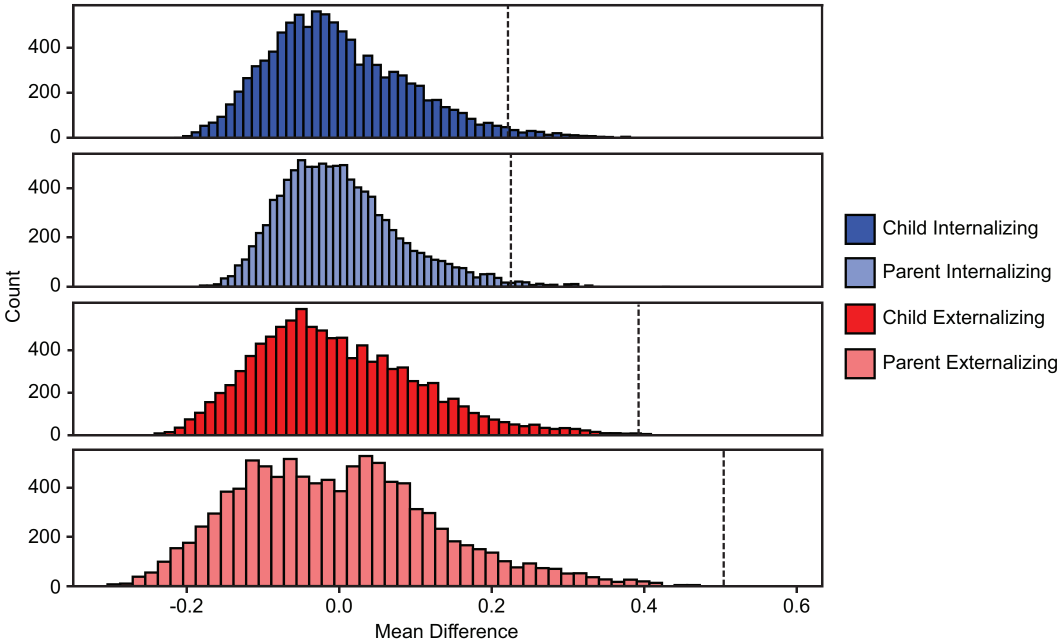


**Supplementary Figure 3**. Correlations of predictive feature weights computed from KRR model outputs were significantly stronger across behavioral measures within the same category than between different categories. Differences between within- and between-category mean correlations for child and parent internalizing and externalizing categories were significantly greater than the null distributions (FDR *q*s ≤0.022). Correlation values were converted to z-scores using Fisher's r-to-z transformation prior to averaging. Histograms display null distributions of mean differences generated through 10000 permutations with shuffled behavioral labels. Dashed lines represent observed mean differences for each of the four categories.

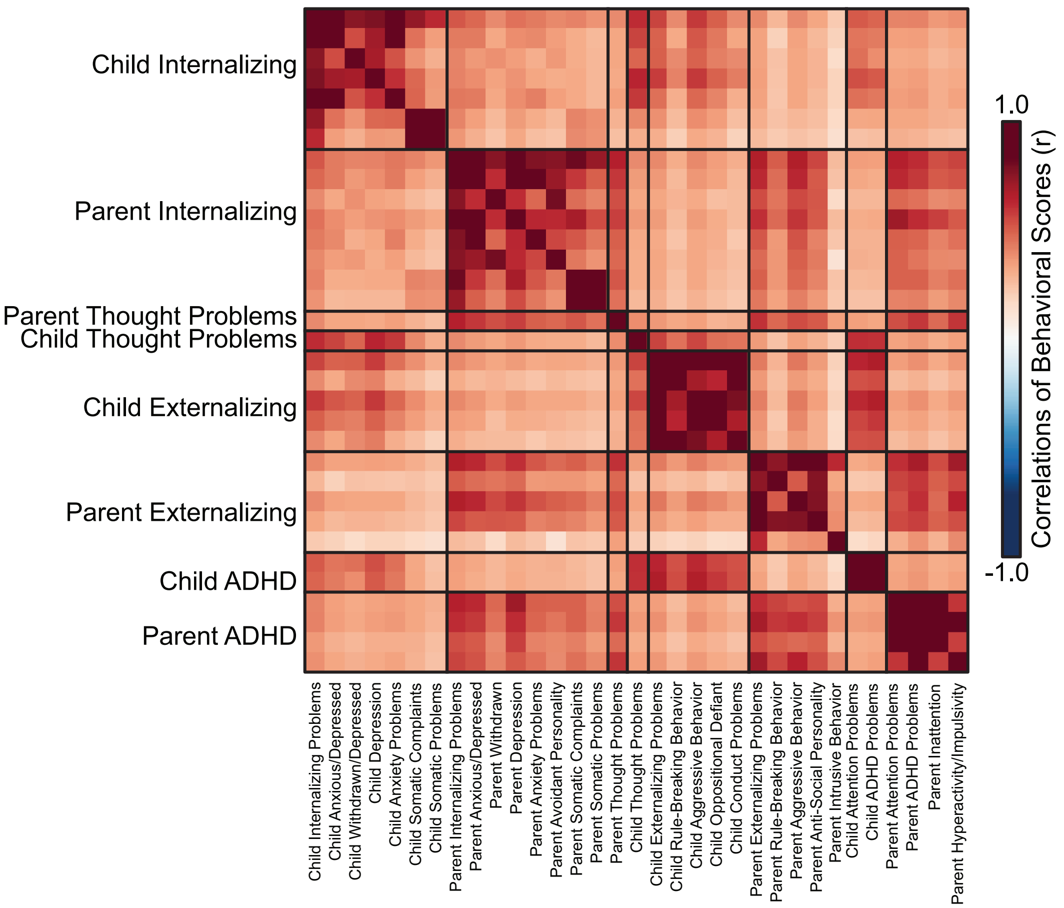


**Supplementary Figure 4.** Pearson's correlation *r* between all pairs of behavioral measures, including measures which failed to achieve better-than-chance brain-behavior predictive accuracy, in the ABCD study. Behavioral measures from the same behavioral categories are grouped together. Warmed colors indicate stronger positive correlations between a pair of behavioral measures. ADHD = Attention-Deficit/Hyperactivity Disorder

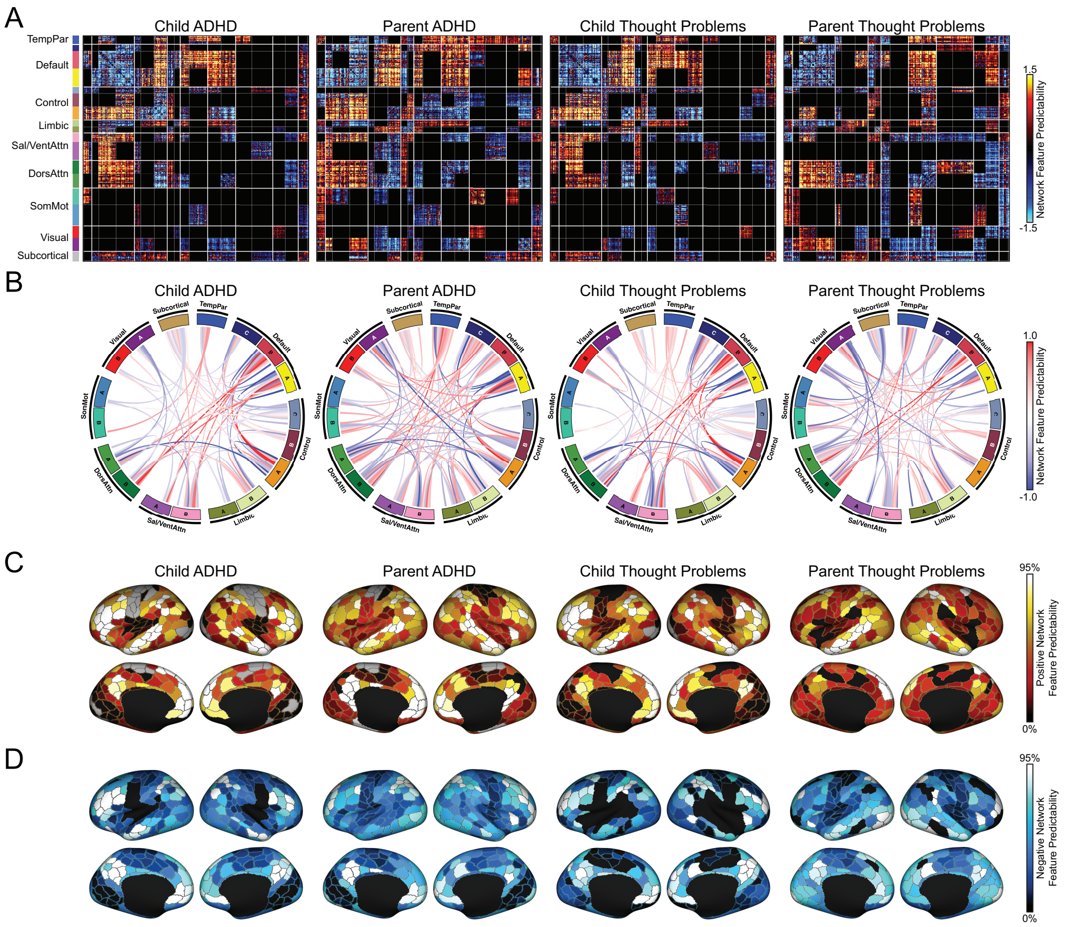


**Supplementary Figure 5.** Shared and unique functional network features predict thought and attention-deficit/hyperactivity disorder (ADHD) problems in children and their parents. (A) Predictive-feature matrices averaged across all behavioral measures within child and parental thought problem and ADHD problem categories and across all brain states. Only weights that are statistically significant and that exhibit the same directionality across all brain states are plotted. Each row corresponds to one cortical parcel, and each matrix entry represents predictive weight associated with functional connectivity between two cortical parcels. All predictive-feature weights were divided by its standard deviation for visualization purposes. (B) Predictive-feature weights averaged within each between-network and within-network block in panel (A). (C) Positive predictive-feature weights summed across rows of panel (A) for each cortical parcel. A more positive value indicates that stronger functional connectivity associated with a given cortical parcel predicts higher behavioral scores in a behavioral category. (D) Negative predictive-feature weights summed across rows of panel (A) for each cortical parcel. A more negative value indicates that weaker functional connectivity associated with a given cortical parcel predicts higher behavioral scores in a behavioral category. In both panels (C) and (D), the color of each cortical parcel indicates the percentile of predictive-feature weights among 400 parcels. (E) The 2D grid displays the proportion of network blocks that exhibit the same directionality across each pair of child behavioral categories relative to the behavioral category represented by each column. Here, each within- and between-network block was coded as 1, 0 or -1 depending on whether sum of predictive-feature weights within that block is greater than, equal to or lesser than 0, resulting in an 18×18 matrix for each behavioral category. The number of network blocks having the same non-zero entries across both matrices associated with each pair of behavioral categories was counted and divided by the total number of non-zero significantly predictive network blocks.

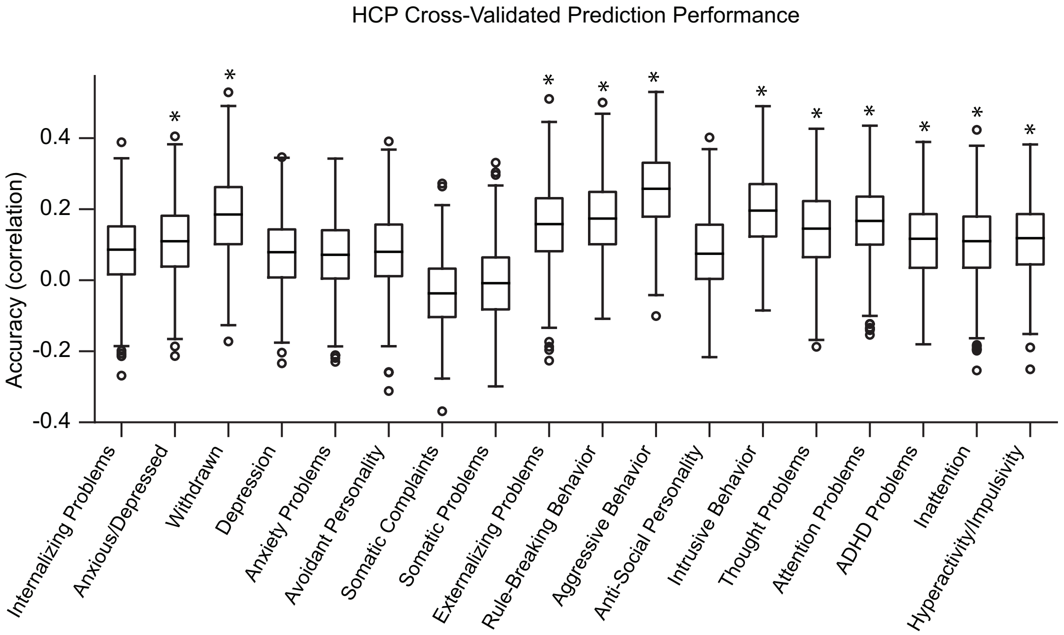


**Supplementary Figure 6.** Cross-validated prediction performance using the KRR model applied to resting-state functional connectivity matrices in HCP subjects. Prediction performance is calculated as mean Pearson's correlation between observed and predicted values across 60 random initiations of 10 cross-validation folds for each behavioral measure selected from HCP dataset. For each boxplot, the top and bottom edges represent upper and lower quartiles of correlation coefficient *r* distribution, and the horizontal line marks the median. Outliers are plotted as circles and are defined as data points outside of the interquartile range. The whiskers extend to the most extreme data points not considered outliers. *s denote above-chance significance after correction for multiple comparisons (FDR $q$<0.05).

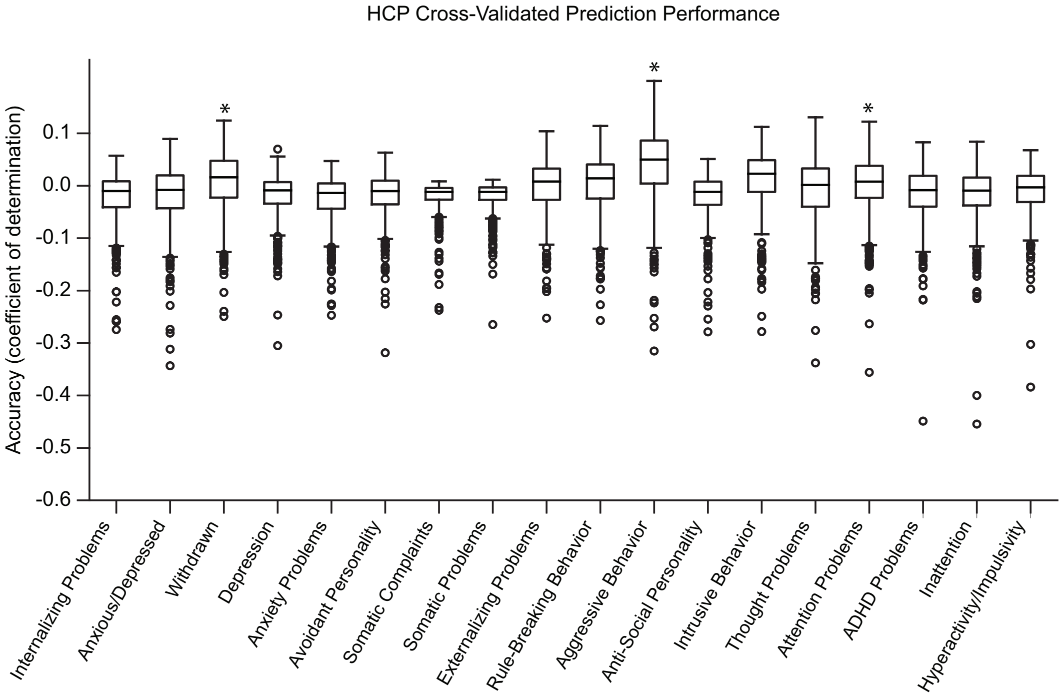


**Supplementary Figure 7.** Cross-validated prediction performance using the KRR model applied to resting-state functional connectivity matrices in HCP subjects. Prediction performance is calculated as mean COD across 60 random initiations of 10 cross-validation folds for each behavioral measure selected from HCP dataset. For each boxplot, the top and bottom edges represent upper and lower quartiles of COD distribution, and the horizontal line marks the median. Outliers are plotted as circles and are defined as data points outside of the interquartile range. The whiskers extend to the most extreme data points not considered outliers. *s denote above-chance significance after correction for multiple comparisons (FDR $q<0.05$).

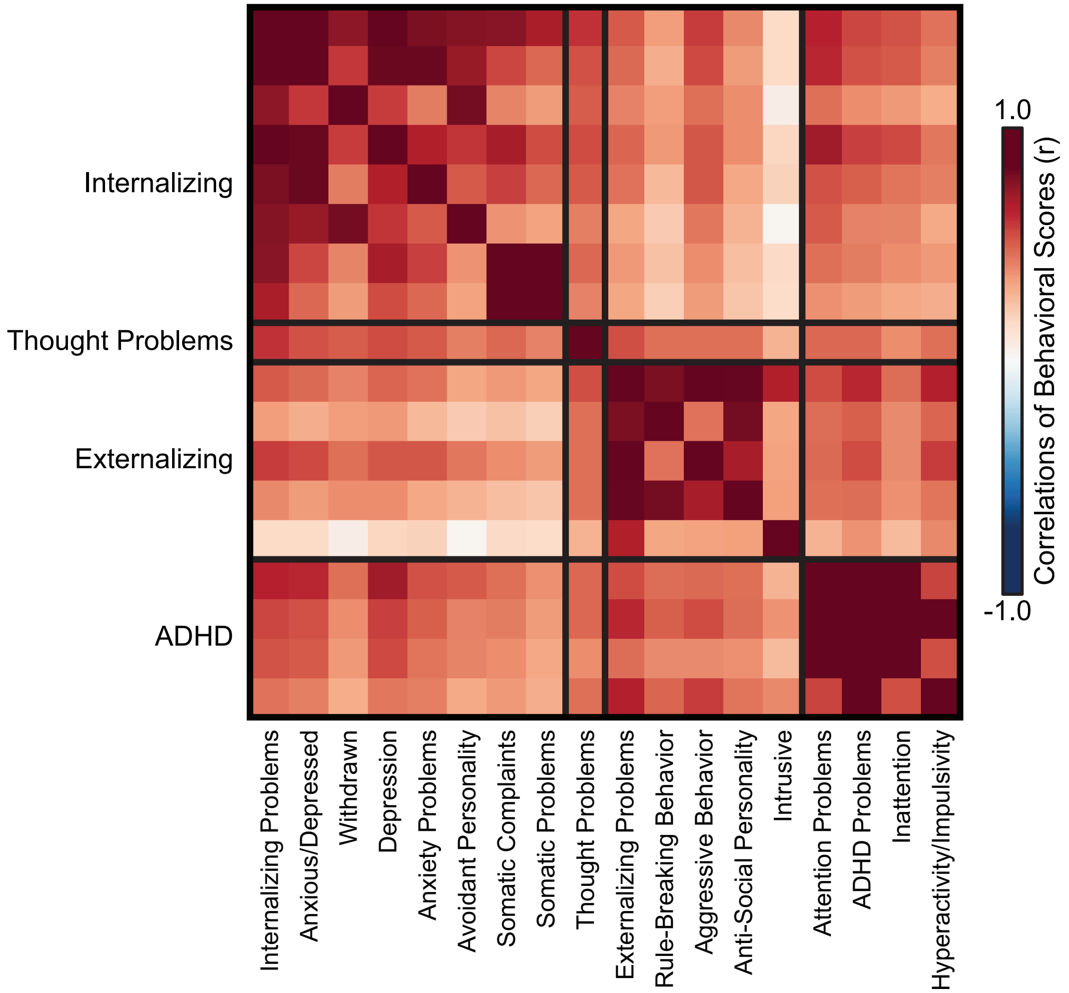


**Supplementary Figure 8.** Pearson's correlation *r* between all pairs of behavioral measures, including measures which failed to achieve better-than-chance brain-behavior predictive accuracy, in the HCP study. Behavioral measures from the same behavioral categories are grouped together. Warmed colors indicate stronger positive correlations between a pair of behavioral measures.

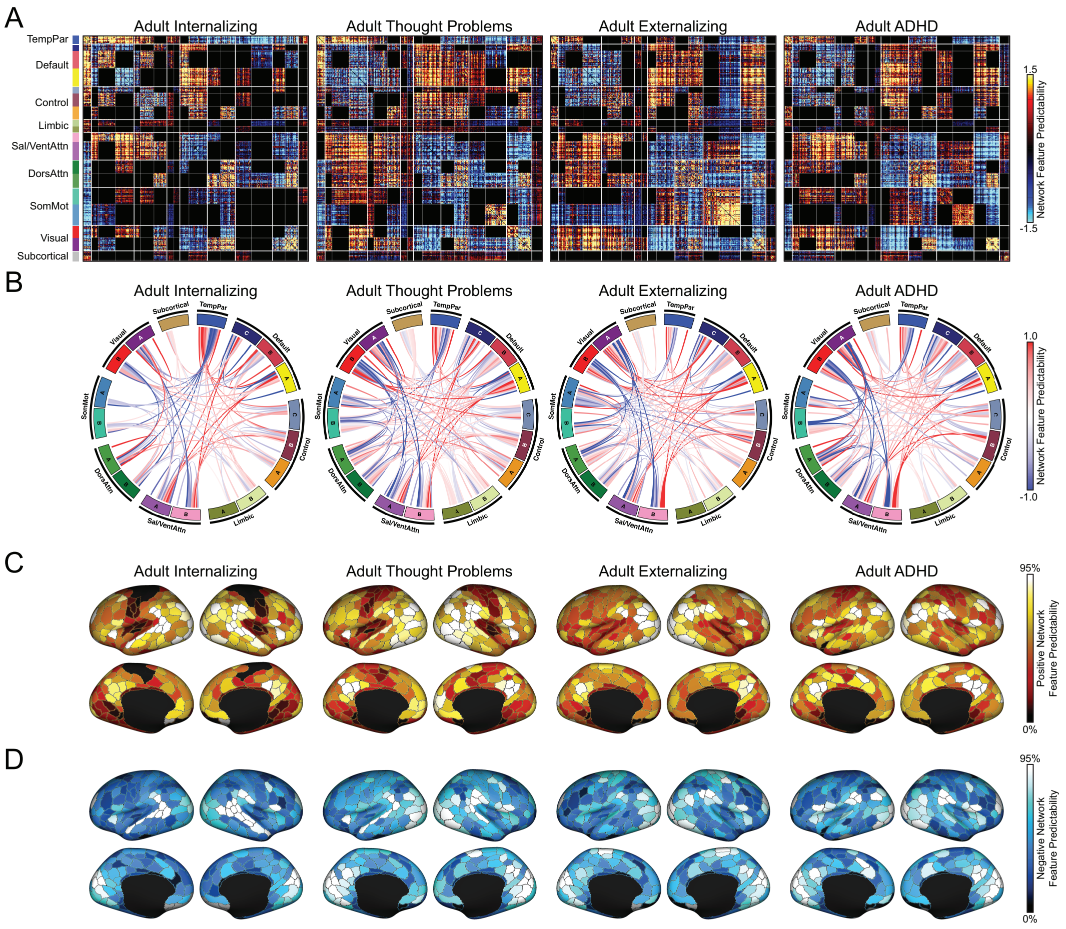


**Supplementary Figure 9.** Similar network features predict internalizing, externalizing, thought and ADHD problems in HCP adults. (A) Predictive-feature matrices averaged across all behavioral measures within each of the four behavioral categories and across all brain states. Graphing conventions are similar to Supplementary Figure 2. (B) Predictive-feature weights averaged within each between-network and within-network block in panel (A). (C) Positive predictive-feature weights summed across rows of panel (A) for each cortical parcel. A more positive value indicates that stronger functional connectivity associated with a given cortical parcel predicts higher behavioral scores in a behavioral category. (D) Negative predictive-feature weights summed across rows of panel (A) for each cortical parcel. A more negative value indicates that weaker functional connectivity associated with a given cortical parcel predicts higher behavioral scores in a behavioral category. In both panels (C) and (D), the color of each cortical parcel indicates the percentile of predictive-feature weights among 400 parcels.

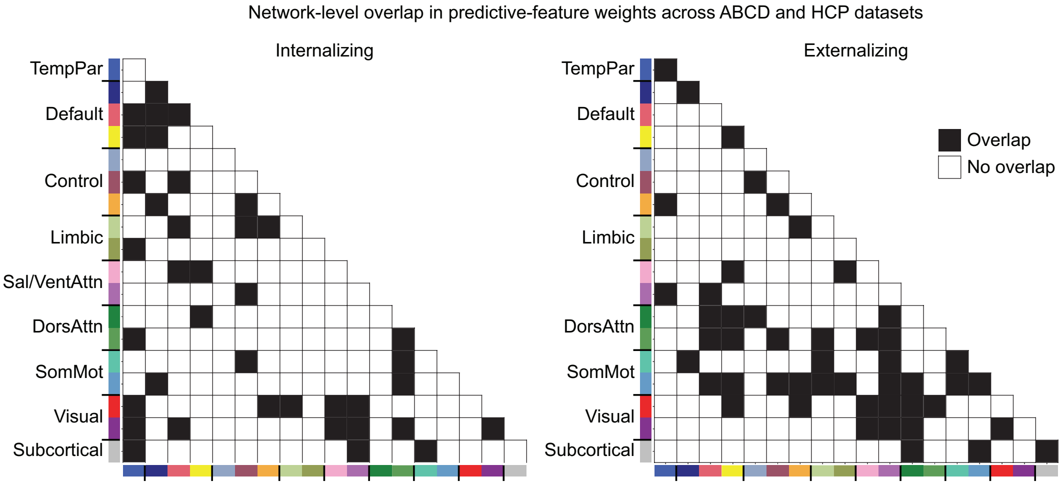


**Supplementary Figure 10.** The matrices display the network blocks which exhibit the same directionality across children and adult internalizing (left) and externalizing (right) categories in black, meaning that they have either positive or negative sums of predictive-feature weights across both categories.

**Supplementary Table S1.** Behavioral categories and associated ABCD study measures in children.

| Behavioral category | Measures (Subscale) |
|---|---|
| Child Internalizing | Child Internalizing Problems (Syndrome Subscale)<br>Child Anxious/Depressed (Syndrome Subscale)<br>Child Withdrawn/Depressed (Syndrome Subscale)<br>Child Depression (DSM-Oriented Subscale)<br>Child Somatic Complaints (Syndrome Subscale)<br>Child Somatic Problems (DSM-Oriented Subscale)<br>Child Anxiety Problems (DSM-Oriented Subscale) |
| Child Externalizing | Child Externalizing Problems (Syndrome Subscale)<br>Child Rule-breaking Behavior (Syndrome Subscale)<br>Child Aggressive Behavior (Syndrome Subscale)<br>Child Oppositional Defiant (DSM-Oriented Subscale)<br>Child Conduct Problems (DSM-Oriented Subscale) |
| Child Thought Problems | Child Thought Problems (Syndrome Subscale) |
| Child ADHD Problems | Child Attention Problems (Syndrome Subscale)<br>Child ADHD Problems (DSM-Oriented Subscale) |
| Parent Internalizing | Parent Internalizing Problems (Syndrome Subscale)<br>Parent Anxious/Depressed (Syndrome Subscale)<br>Parent Withdrawn (Syndrome Subscale)<br>Parent Depression (DSM-Oriented Subscale)<br>Parent Somatic Complaints (Syndrome Subscale)<br>Parent Somatic Problems (DSM-Oriented Subscale)<br>Parent Anxiety Problems (DSM-Oriented Subscale)<br>Parent Avoidant Personality Problems (DSM-Oriented Subscale) |
| Parent Externalizing | Parent Externalizing Problems (Syndrome Subscale)<br>Parent Rule-breaking Behavior (Syndrome Subscale)<br>Parent Aggressive Behavior (Syndrome Subscale)<br>Parent Intrusive Behavior (Syndrome Subscale)<br>Parent Antisocial Personality Problems (DSM-Oriented Subscale) |
| Parent Thought Problems | Parent Thought Problems (Syndrome Subscale) |
| Parent ADHD Problems | Parent Attention Problems (Syndrome Subscale)<br>Parent ADHD Problems (DSM-Oriented Subscale)<br>Parent Inattention (DSM-Oriented Subscale)<br>Parent Hyperactivity-Impulsivity (DSM-Oriented Subscale) |

**Note**: We selected 25 clinical assessment dimensions related to internalizing and externalizing problems from all mental health assessments in the Adolescent Brain Cognitive Development

(ABCD) study[1]. Specifically, we included scores from the Achenbach Child Behavior Check List (CBCL)[2] and the Achenbach Adult Self-Report (ASR) scales[3]. ABCD = Adolescent Brain Cognitive Development; DSM = Diagnostic and Statistical Manual of Mental Disorders; ADHD = Attention-Deficit/Hyperactivity Disorder.

**Supplementary Table S2.** Behavioral categories and associated HCP study measures in adults.

| Behavioral category | Measures (Subscale) |
|---|---|
| Adult Internalizing | Internalizing Problems (Syndrome Subscale)<br>Anxious/Depressed (Syndrome Subscale)<br>Withdrawn (Syndrome Subscale)<br>Depression (DSM-Oriented Subscale)<br>Somatic Complaints (Syndrome Subscale)<br>Somatic Problems (DSM-Oriented Subscale)<br>Anxiety Problems (DSM-Oriented Subscale)<br>Avoidant Personality Problems (DSM-Oriented Subscale) |
| Adult Externalizing | Externalizing Problems (Syndrome Subscale)<br>Rule-breaking Behavior (Syndrome Subscale)<br>Aggressive Behavior (Syndrome Subscale)<br>Intrusive Behavior (Syndrome Subscale)<br>Antisocial Personality Problems (DSM-Oriented Subscale) |
| Adult Thought Problems | Thought Problems (Syndrome Subscale) |
| Adult ADHD Problems | Attention Problems (Syndrome Subscale)<br>ADHD Problems (DSM-Oriented Subscale)<br>Inattention (DSM-Oriented Subscale)<br>Hyperactivity/Impulsivity (DSM-Oriented Subscale) |

**Note***:* We selected 18 clinical assessment dimensions related to internalizing and externalizing problems from all mental health assessments in the Human Connectome Project (HCP) study[4]. Specifically, we included scores from the Achenbach Adult Self-Report (ASR) scales[3]. HCP = Human Connectome Project; DSM = Diagnostic and Statistical Manual of Mental Disorders; ADHD = Attention-Deficit/Hyperactivity Disorder

**Supplementary Table S3.** Table with the original ABCD variable names corresponding to the measures used in this study

| Measure | Original ABCD Variable Name | ABCD Data File |
|---|---|---|
| Child Internalizing Problems | cbcl_scr_syn_internal_r | abcd_cbcls01.txt |
| Child Anxious/Depressed | cbcl_scr_syn_anxdep_r | abcd_cbcls01.txt |
| Child Withdrawn/Depressed | cbcl_scr_syn_withdep_r | abcd_cbcls01.txt |
| Child Depression | cbcl_scr_dsm5_depress_r | abcd_cbcls01.txt |
| Child Somatic Complaints | cbcl_scr_syn_somatic_r | abcd_cbcls01.txt |
| Child Somatic Problems | cbcl_scr_dsm5_somaticpr_r | abcd_cbcls01.txt |
| Child Anxiety Problems | cbcl_scr_dsm5_anxdisord_r | abcd_cbcls01.txt |
| Child Externalizing Problems | cbcl_scr_syn_external_r | abcd_cbcls01.txt |
| Child Rule-breaking Behavior | cbcl_scr_syn_rulebreak_r | abcd_cbcls01.txt |
| Child Aggressive Behavior | cbcl_scr_syn_aggressive_r | abcd_cbcls01.txt |
| Child Oppositional Defiant | cbcl_scr_dsm5_opposit_r | abcd_cbcls01.txt |
| Child Conduct Problems | cbcl_scr_dsm5_conduct_r | abcd_cbcls01.txt |
| Child Thought Problems | cbcl_scr_syn_thought_r | abcd_cbcls01.txt |
| Child Attention Problems | cbcl_scr_syn_attention_r | abcd_cbcls01.txt |
| Child ADHD Problems | cbcl_scr_dsm5_adhd_r | abcd_cbcls01.txt |
| Parent Internalizing Problems | asr_scr_internal_r | abcd_asrs01.txt |
| Parent Anxious/Depressed | asr_scr_anxdep_r | abcd_asrs01.txt |
| Parent Withdrawn | asr_scr_withdrawn_r | abcd_asrs01.txt |
| Parent Depression | asr_scr_depress_r | abcd_asrs01.txt |
| Parent Somatic Complaints | asr_scr_somatic_r | abcd_asrs01.txt |
| Parent Somatic Problems | asr_scr_somaticpr_r | abcd_asrs01.txt |
| Parent Anxiety Problems | asr_scr_anxdisord_r | abcd_asrs01.txt |
| Parent Avoidant Personality Problems | asr_scr_avoidant_r | abcd_asrs01.txt |
| Parent Externalizing Problems | asr_scr_external_r | abcd_asrs01.txt |
| Parent Rule-breaking Behavior | asr_scr_rulebreak_r | abcd_asrs01.txt |
| Parent Aggressive Behavior | asr_scr_aggressive_r | abcd_asrs01.txt |
| Parent Intrusive Behavior | asr_scr_intrusive_r | abcd_asrs01.txt |
| Parent Antisocial Personality Problems | asr_scr_antisocial_r | abcd_asrs01.txt |

| | | |
|---|---|---|
| Parent Thought Problems | asr_scr_thought_r | abcd_asrs01.txt |
| Parent Attention Problems | asr_scr_attention_r | abcd_asrs01.txt |
| Parent ADHD Problems | asr_scr_adhd_r | abcd_asrs01.txt |
| Parent Inattention | asr_scr_inattention_r | abcd_asrs01.txt |
| Parent Hyperactivity-Impulsivity | asr_scr_hyperactivity_r | abcd_asrs01.txt |

**Supplementary Table S4**. Table with the original HCP variable names corresponding to the measures used in this study

| Measure | Original HCP Variable Name |
|---|---|
| Internalizing Problems | ASR_Intn_Raw |
| Anxious/Depressed | ASR_Anxd_Raw |
| Withdrawn | ASR_Witd_Raw |
| Depression | DSM_Depr_Raw |
| Anxiety Problems | DSM_Anxi_Raw |
| Avoidant Personality Problems | DSM_Avoid_Raw |
| Somatic Complaints | ASR_Soma_Raw |
| Somatic Problems | DSM_Somp_Raw |
| Thought Problems | ASR_Thot_Raw |
| Externalizing Problems | ASR_Extn_Raw |
| Rule-breaking Behavior | ASR_Rule_Raw |
| Aggressive Behavior | ASR_Aggr_Raw |
| Antisocial Personality Problems | DSM_Antis_Raw |
| Intrusive Behavior | ASR_Intr_Raw |
| Attention Problems | ASR_Attn_Raw |
| ADHD Problems | DSM_Adh_Raw |
| Inattention | DSM_Inat_Raw |
| Hyperactivity/Impulsivity | DSM_Hype_Raw |

**Supplementary Table S5**. Distribution of ABCD analytical sample (n=2262) by site, scanner and their assigned site cluster

| ABCD Site | Make | Model | N | Site Cluster |
|---|---|---|---|---|
| 16 | Siemens | Prisma | 337 | A |
| 13 | GE | Discovery MR750 | 192 | B |
| 4 | GE | Discovery MR750 | 170 | C |
| 22 | GE | Discovery MR750 | 13 | C |
| 14 | Siemens | Prisma/Prisma fit | 165 | D |
| 15 | Siemens | Prisma fit | 37 | D |
| 6 | Siemens | Prisma fit | 150 | E |
| 9 | Siemens | Prisma fit | 66 | E |
| 10 | GE | Discovery MR750 | 161 | F |
| 11 | Siemens | Prisma | 71 | F |
| 3 | Siemens | Prisma | 160 | G |
| 5 | Siemens | Prisma fit | 72 | G |
| 2 | Siemens | Prisma fit | 129 | H |
| 7 | Siemens | Prisma fit | 71 | H |
| 8 | GE | Discovery MR750 | 74 | I |
| 20 | Siemens | Prisma/Prisma fit | 103 | I |
| 12 | Siemens | Prisma fit | 116 | J |
| 18 | GE | Discovery MR750 | 82 | J |
| 21 | Siemens | Prisma fit/Prisma | 93 | J |

**Supplementary Table S6**. Demographic information for included and excluded participants from ABCD study release 2.0.1

| | Included | Excluded |
|---|---|---|
| N | 2262 | 9613 |
| Age in months, mean (± SD) | 120.15 (7.48) | 118.66 (7.43) |
| Sex, (% male) | 1030 (45.53) | 5158 (53.56) |
| Race/ethnicity (%) | | |
| Caucasian | 1335(59.02) | 4839 (50.24) |
| African American | 201 (8.89) | 1578 (16.38) |
| Hispanic | 425 (18.79) | 1982 (20.58) |
| Asian | 60 (2.65) | 192 (1.99) |
| Other | 236 (10.43) | 1009 (10.48) |

**Supplementary Table S7**. Demographic information for included and excluded participants from HCP S1200 Data Release

| | Included | Excluded |
|---|---|---|
| N | 752 | 454 |
| Age in years, mean (± SD) | 28.61 (3.72) | 29.21 (3.62) |
| Sex, (% male) | 353 (46.94) | 197 (43.39) |
| Race/ethnicity (%) | | |
| Caucasian | 584 (77.66) | 303 (66.74) |
| African American | 89 (11.84) | 104 (22.91) |
| Asian/Hawaiian National/Other Pacific Islander | 45 (5.98) | 24 (5.29) |
| Other/Unreported | 14 (1.86) | 11 (2.42) |

**Supplemental Method S1. Single-Kernel Ridge Regression**

The following section is adapted from Kong et al., 2019[5]. Suppose there are M training subjects in the HCP dataset. Let $y_m$ be the behavioral measure for the *m*-th training subject and $FC_m$ be the vectorized resting-state functional connectivity (FC) matrix (consider only the lower triangle) for the *m*-th training subject. The single-kernel ridge regression model can be written as:

$$y_m = \beta_0 + \sum_{j=1}^{M} \alpha_j K(FC_j, FC_m) \; \boldsymbol{s.t.} \; m, j \in \{1, 2, \ldots, \mathrm{M}\}$$

where the $\beta_0$ is the bias term, $K(FC_j, FC_m)$ is the similarity of the FC vectors between the *j*-th training subject and the *m*-th training subject. We used correlation as the similarity measure as motivated by previous fingerprinting and behavioral prediction studies [6–8]. As such, $K(FC_j, FC_m)$ is the functional connectivity similarity between the *j*-th and *m*-th training subject, and $\alpha_j$ is the alpha weight for the *j*-th training subject. Let $\boldsymbol{y} = [y_1, y_2, \cdots, y_M]^T$, $\boldsymbol{\alpha} = [\alpha_1, \alpha_2, \cdots, \alpha_M]^T$ and $\mathbb{K}$ be the M×M correlation matrix across all pairs of training subjects where the (*j, i*)-th entry is $K(FC_j, FC_m)$. We estimated optimal $\beta_0$ and $\boldsymbol{\alpha}$ by minimizing the loss function below:

$$\underset{(\beta_0, \boldsymbol{\alpha})}{\operatorname{argmin}} \frac{1}{2}(\boldsymbol{y} - \beta_0 - \mathbb{K}\boldsymbol{\alpha})^T(\boldsymbol{y} - \beta_0 - \mathbb{K}\boldsymbol{\alpha}) + \frac{\lambda}{2}\boldsymbol{\alpha}^T \mathbb{K}\boldsymbol{\alpha}$$

Here, $\lambda$ controls the importance of the $l_2$ regularization term and was optimized by inner-loop cross-validation within the training set. The regularization and model parameters ($\lambda$, $\boldsymbol{\alpha}$ and $\beta_0$) estimated from the training set were then used to predict behavior of the *t*-th test subject by

$$\hat{y}_t = \beta_0 + \sum_{m=1}^{M} \alpha_m K(FC_m, FC_t)$$

**Supplemental Method S2. Multi-Kernel Ridge Regression**

Suppose there are M training subjects and R brain states in the ABCD dataset. Let $y_m$ and $FC_{mr}$ be the behavioral measure for the *m*-th training subject and the vectorized FC matrix (consider only the lower triangle) for the *m*-th training subject for the *r*-th brain state respectively. The multiKRR model can be written as:

$$y_m = \beta_0 + \sum_{r=1}^{R}\sum_{j=1}^{M} \alpha_{jr} K(FC_{jr}, FC_{mr}) \ \boldsymbol{s.t.} \ m, j \in \{1, 2, \dots, \mathrm{M}\} \ and \ r \in \{1, 2, \dots, \mathrm{R}\}$$

or

$$\boldsymbol{y_{train}} = \beta_0 + \sum_{r=1}^{R} \boldsymbol{\alpha_r} \mathbb{K}_{\boldsymbol{r}}$$

where the $\beta_0$ is the bias term, $K(FC_{jr}, FC_{mr})$ is defined as correlation of the FC vectors between the *j*-th training subject and the *m*-th training subject for the *r*-th brain state. As such, $\mathbb{K}_r$ is an M×M correlation matrix across all pairs of training subjects for brain state r where the (*j, i*)-th entry is $K(FC_{jr}, FC_{mr})$. $\boldsymbol{y}_{train}$ and $\boldsymbol{\alpha}_r$ are behavioral measures and alpha weights for the *r*-th brain state across all training subjects. We estimated optimal $\beta_0$ and $\boldsymbol{\alpha}_r$ by minimizing the loss function below using Gaussian-process optimization (Kawaguchi et al., 2015):

$$\underset{(\beta_0, \boldsymbol{\alpha})}{\operatorname{argmin}} \frac{1}{2}\left(\boldsymbol{y} - \beta_0 - \sum_{r=1}^{R} \mathbb{K}_{\boldsymbol{r}} \boldsymbol{\alpha}_r\right)^T \left(\boldsymbol{y} - \beta_0 - \sum_{r=1}^{R} \mathbb{K}_{\boldsymbol{r}} \boldsymbol{\alpha}_r\right) + \frac{\lambda_r}{2}\sum_{r=1}^{R} \boldsymbol{\alpha}_r{}^T \mathbb{K}_{\boldsymbol{r}} \boldsymbol{\alpha}_r$$

Here, $\lambda_r$ controls the weighting of the $l_2$ regularization term for the *r*-th brain state and was optimized by inner-loop cross-validation within the training set. The regularization and model parameters ($\lambda_r$, $\boldsymbol{\alpha}_r$ and $\beta_0$) estimated from the training set were then used to predict behavior of the *t*-th test subject by

$$\hat{y}_t = \beta_0 + \sum_{r=1}^{R}\sum_{m=1}^{M} \alpha_{mr} K(FC_{mr}, FC_{tr})$$

**Supplemental Method S3. Coefficients of Determination**

Suppose that there are N test subjects, the predictive accuracy of KRR and multiKRR models is measured by coefficients of determination (COD) which determines the proportion of variance explained by fitting each model,

$$COD = 1 - \frac{\sum_{t=1}^{N} (\hat{y}_t - y_t)^2}{\sum_{t=1}^{N} (\bar{y}_{train} - y_t)^2}$$

where $y_t$ and $\hat{y}_t$ are the ground truth and the predicted behavior measure of the *t*-th test subject, and $\bar{y}_{train}$ is the mean behavior dimension across all training subjects. A larger COD indicates more accurate prediction, while a negative COD means that the mean behavior dimension across the training set is a better predictor than the model.
